# mRNA-based prime-and-pull vaccination combines benefits of subcutaneous and mucosal BCG vaccination

**DOI:** 10.64898/2026.07.21.739464

**Authors:** Ana Maria Valencia-Hernandez, Guangzu Zhao, Julia Seifert, Harindra D. Sathkumara, Socorro Miranda-Hernadez, Munish Puri, Andreas Kupz

**Affiliations:** Australian Institute of Tropical Health and Medicine, James Cook University, Cairns & Townsville, QLD, Australia

**Keywords:** Tuberculosis, vaccine development, mRNA vaccine, mucosal vaccination, BCG, prime-and-pull

## Abstract

Pulmonary vaccination has been proposed as a strategy to improve protection against tuberculosis, due to the generation of immune cells that more efficiently survey and eliminate infected cells within the lung. However, uncontrolled replication and excessive inflammation associated with mucosal delivery of live-attenuated vaccines highlight the need for alternative strategies that balance efficacy and safety. Here, we describe a prime-and-pull vaccination approach in which systemic immunity is established by subcutaneous BCG vaccination, followed by the induction of local lung immunity through mucosal delivery of lipid nanoparticle–formulated multi-antigen mRNA. Proof-of-concept studies using a model antigen demonstrated the induction of polyfunctional antigen-specific T cells in the lung with minimal inflammatory cell infiltration, compared with mucosal BCG vaccination. Eight *Mycobacterium tuberculosis*- and BCG-derived proteins were subsequently selected to generate four multi-antigen mRNA constructs. *In vivo* vaccination and challenge experiments demonstrated that this prime-and-pull strategy is well tolerated and confers protective immunity against tuberculosis in a murine model. Collectively, these data support a modular mRNA-based prime-and-pull vaccination strategy as a translational approach that bridges the improved immunogenicity and efficacy of mucosal BCG vaccination with the safety profile of parenteral BCG administration.

**Graphical Abstract:** 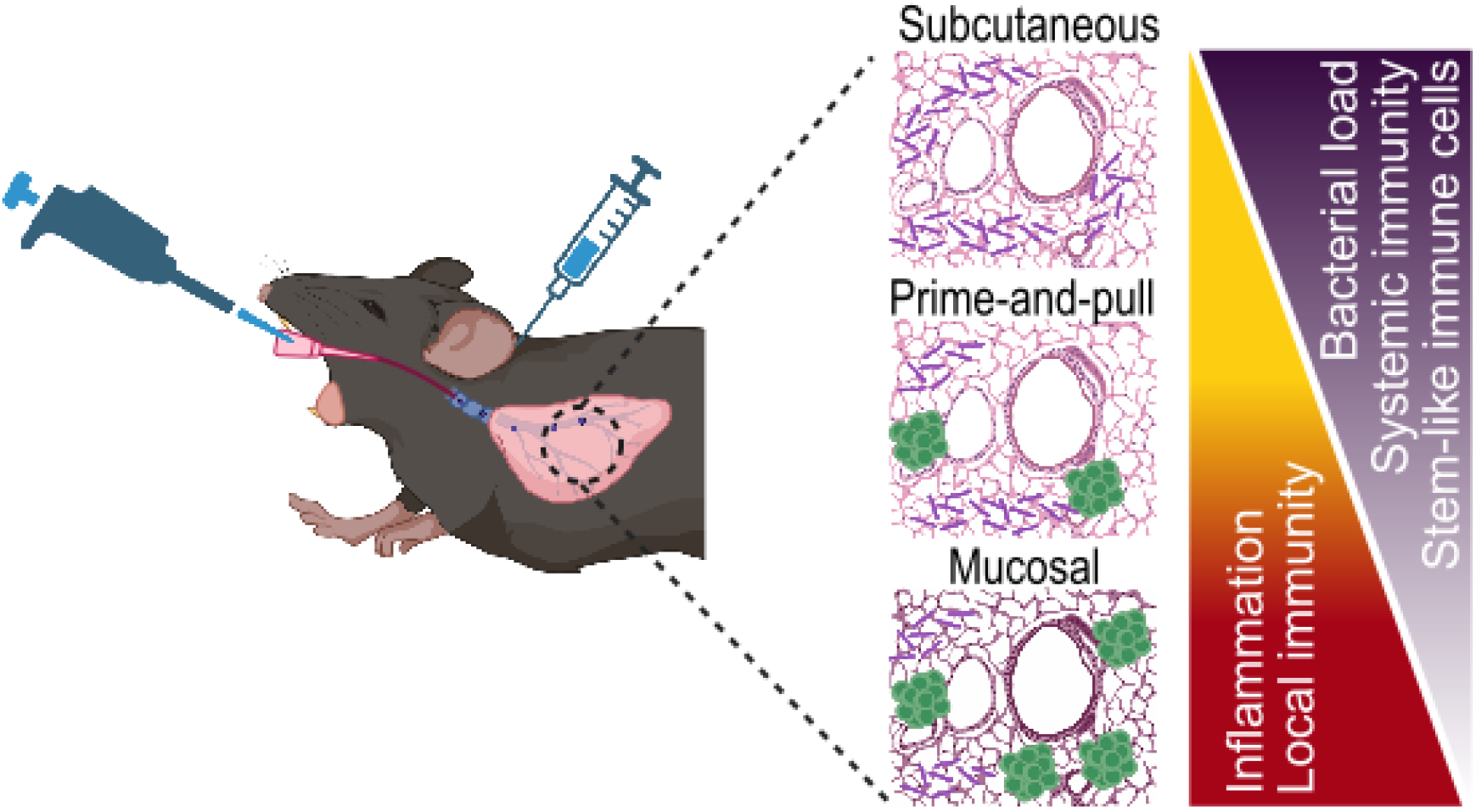

## Introduction

The only licensed tuberculosis (TB) vaccine, *Mycobacterium bovis* Bacillus Calmette-Guérin (BCG), protects children against severe forms of disease, reduces all-cause infant mortality, and confers heterologous protection against multiple infectious diseases beyond childhood^1–5^. However, protection against pulmonary TB (the most prevalent and transmissible form of the disease) varies widely, ranging from >70% in Nordic countries to <20% in TB-endemic regions^6–11^. As a result, there is an urgent need for more effective TB vaccination strategies, particularly in settings where BCG efficacy is lowest. Although several vaccine candidates have advanced through clinical development over recent decades, none have surpassed BCG in protective efficacy, which remains the benchmark for TB vaccination^12^.

Mucosal vaccination has emerged as an attractive approach due to its potential to induce superior protection compared with conventional parenteral delivery. Protective mucosal immunity relies on the generation of robust local immune responses at sites of pathogen entry and is associated with the formation of tissue-resident memory (T_RM_) cells^13–15^. However, mucosal vaccination faces several biological barriers, including the presence of mucus, antimicrobial factors, and tolerogenic environments that can limit antigen delivery and immune priming^16^. Effective mucosal immunization therefore requires a finely tuned inflammatory response that promotes immunity while preserving tissue integrity and function. In the respiratory tract, immune cells, including T_RM_ populations, are primed and maintained within transient inflammatory niches induced by microbial exposure or other inflammatory stimuli^17, 18^. A further challenge is the comparatively short lifespan of lung T_RM_ cells relative to other tissues^19–21^. Their rapid decline has been attributed to cell death and redistribution to the mediastinal lymph node (medLN), potentially reflecting an intrinsic incompatibility between sustained inflammation and efficient gas exchange in the lung^21, 22^.

To date, only a limited number of mucosal delivered TB vaccine candidates have entered clinical evaluation. These include viral-vectored vaccines expressing *Mycobacterium tuberculosis* (*Mtb*) antigens, such as MVA85A, AdHu5Ag85A, ChAdOx1-85A, and TB/FLU-05E, which have been assessed for safety following aerosol or intranasal administration in Phase I trials^23–27^. Among live-attenuated vaccines (LAVs), recent studies have investigated aerosolized BCG as a controlled challenge model in vaccinated and unvaccinated individuals (NCT02709278, NCT06670755, NCT06246851, NCT03912207, NCT04777721)^28^. Although high-dose aerosol BCG was generally well tolerated, with only a trend toward increased adverse events^28^, concerns remain regarding potentially uncontrolled replication and excessive inflammation following mucosal delivery of LAVs, underscoring the need for alternative strategies.

To address these challenges, we sought to develop an mRNA-based prime-and-pull vaccination strategy designed to preserve immunogenicity while limiting inflammation and avoiding the risk of microbial replication. We hypothesised that subcutaneous BCG priming would establish systemic immunity, followed by mucosal administration of lipid nanoparticle (LNP)-formulated multi-antigen mRNA to promote local lung immunity. Mucosal delivery of mRNA should enable transient antigen expression within the lung, facilitating the recruitment and differentiation of antigen-specific T cells into T_RM_ cell populations capable of responding to *Mtb*-infected cells. Because mRNA vaccines are non-replicating, this approach was expected to reduce the inflammatory burden associated with live mucosal vaccination.

We first evaluated whether customized LNPs could deliver mRNA to lung cells and support local protein expression in vivo. Using the well-characterized, immunodominant *Mtb*-derived 6 kDa early secretory antigenic target (ESAT-6) as a model antigen, we then showed that a single high-dose prime-and-pull regimen, rather than multiple low-dose administrations, induced robust populations of polyfunctional antigen-specific T cells in the lungs of vaccinated mice, with minimal cellular infiltration relative to mucosal BCG vaccination. Guided by these findings, we next selected eight *Mtb*- and BCG-derived antigens to generate four multi-antigen mRNA constructs formulated in customized LNPs as a proof-of-principle evaluation of a broader multivalent strategy. *In vivo* vaccination and challenge studies showed that this prime-and-pull approach was well tolerated in mice and supported protective immunity in a murine model. Collectively, these data provide proof of principle for a vaccination strategy that seeks to combine the immunological advantages of mucosal immune recruitment with the reduced inflammatory burden of non-replicating mRNA delivery.

## Materials and methods

### Bacterial strains and culture conditions

*Mycobacterium bovis* BCG SSI Danish (hereafter referred to as BCG), recombinant BCG expressing the *Mtb* ESX 1 locus (BCG::ESX1*^Mtb^*, also referred to as BCG::RD1), and *Mtb* H37Rv were cultured in Middlebrook 7H9 broth (BD Difco) supplemented with 10% oleic albumin dextrose catalase (OADC; USBiological Life Sciences), 0.05% Tween-80, 0.2% glycerol, and appropriate antibiotics at 37°C until mid-logarithmic phase, harvested, and stored in sterile PBS containing 15% glycerol at −80°C until use.

### mRNA molecules

Commercial reporter mRNA encoding enhanced green fluorescent protein (ARCA-EGFP mRNA, 5-moUTP) was purchased from APExBIO. All custom mRNA constructs were obtained from Messenger Bio (https://www.messenger.bio/mrna). These constructs included N1-methylpseudouridine modifications, Cap1 capping, optimized 5′ and 3′ untranslated regions, and a 100-nt poly(A) tail. Codon optimization for mammalian expression was performed as part of the Messenger Bio service.

For proof-of-concept studies, mRNA encoding ESAT-6 was generated. For subsequent experiments, four multi-antigen mRNA constructs were designed using the general architecture signal peptide (SP)–antigen 1–linker–antigen 2. Coding sequences for the selected antigens were obtained from the Mycobrowser database (https://mycobrowser.epfl.ch).

The multi-antigen constructs were as follows:

- Molecule 1, SP–HBHA–linker–HRP1 (1101 nt).
- Molecule 2, SP–HspX–linker–CFP10 (807 nt).
- Molecule 3, SP–Ag85B–linker–VapB47 (1347 nt).
- Molecule 4, SP–PPE18–linker–PE13 (1542 nt).

The human IgE signal peptide sequence was *atggattggacctggattctgtttctggtggcggcggcgacccgcgtgcatagc*, and the linker sequence was *agggcggcggcggcagc*.

All mRNA constructs were mixed in equal amounts (250 μg each), aliquoted into two portions, and stored at −80°C before formulation.

### Lipid nanoparticle formulation

mRNA constructs were formulated into “stealth” lipid nanoparticles (sLNPs) by the Griffith University Nanomedicine Biofoundry. sLNPs were composed of 1,2-dioleoyl-3-trimethylammonium-propane (DOTAP), dioleoylphosphatidylethanolamine (DOPE), cholesterol, and C16 PEG-ceramide and prepared in an aqueous sucrose solution (92.5 mg/mL). Lyophilized mRNA-sLNPs were stored at −80°C for up to six months. Prior to use, sLNPs were equilibrated to room temperature for 30 min and rehydrated with nuclease-free water (Thermo Fisher Scientific) to a final concentration of 0.2 μg/μL. Samples were gently swirled, incubated at room temperature for 2 h, and stored at 4°C for up to 48 h before *in vivo* administration. Quality control analysis of representative batches indicated a mean particle diameter of ∼200 nm and an encapsulation efficiency of 97%.

### Mice

Female C57BL/6 mice (6–8 weeks old) were obtained from the Animal Resource Centre (Perth, Australia) for GFP and ESAT-6 proof-of-concept experiments. For multi-antigen prime-and-pull studies, female C57BL/6 × BALB/c F1 hybrid mice were bred and maintained in-house. All animals were acclimatized for at least one week prior to experimentation and housed under specific pathogen-free conditions in biosafety level 2 or 3 facilities at the Australian Institute of Tropical Health and Medicine (AITHM), James Cook University. All procedures were conducted in accordance with National Health and Medical Research Council (NHMRC) guidelines and approved by the James Cook University Animal Ethics Committee (approval number A2837).

### Vaccination protocols

Frozen BCG stocks were thawed, washed, sonicated, and diluted in sterile PBS prior to immunization. Mice were primed either subcutaneously (SC; 1 × 10⁶ CFU in 50 μL) or intratracheally (IT; 1 × 10⁵ CFU in 100 μL). For prime-and-pull vaccination, mice received IT administration of 5 or 10 μg mRNA-sLNPs at days 7, 14, or 21 following BCG priming. Control groups received empty LNPs (lacking mRNA) or mRNA-sLNPs alone without prior BCG priming.

### *Mtb* challenge and bacterial burden

Sixty days post-vaccination, mice were challenged with a very low dose of 10–70 CFU *Mtb* H37Rv using a Glas-Col aerosol exposure system. Lung and spleen homogenates were plated on 7H11 agar and incubated for 5–8 weeks at 37°C before CFU enumeration.

### Tissue processing and cell preparation

At weeks 4 or 8 post-BCG immunization, spleen, lung, medLN and BALF were collected. Lungs were processed using a GentleMACS™ Dissociator (Miltenyi Biotec) and digested with collagenase D (7.5 μg/mL), collagenase VIII (1.75 μg/mL), and DNase I (200 μg/mL) all from Sigma for 30 min at 37°C. Cell suspensions were filtered through 70 μm strainers, subjected to erythrocyte lysis, and washed in FACS buffer. Spleens and medLNs were mechanically dissociated and processed similarly. BALF was centrifuged, erythrocytes lysed, and cells washed; supernatants were stored at −80°C for antibody analysis.

### Cell surface and intracellular cytokine staining

Splenocytes, lung and medLN cells were cultured in presence or absence of 2 μL/mL of cell stimulation cocktail (eBioscience, 00-4975-93) and brefeldin A (BD, 555029) in RPMI complete media for 5-6 hours at 37°C in a humidified chamber and 5% CO_2_. Dead cells were excluded by Horizon™ Fixable Viability Stain 780 (BD Biosciences) staining. The cell surface was stained with the following antibodies and fluorochromes: TCRβ (BUV737; clone H57-597), CD8a (AF800; clone 53-6.7), CD4 (BUV395; clone GK1.5), CD44 (BV421; clone IM7), CD69 (PE/CF594; clone H1.2F3), CD103 (BV710; clone M290), CD11c (BUV395; clone HL3), I-A/I-E (BV711; clone M5/114), CD45.2 (BV650; clone 104), CD64 (BV786; clone X54-5/7.1) from BD Horizon; CD62L (PE/Cy7; clone MEL-14), MerTK (BV421; clone 2B10C42), CD3 (BV510; clone 17A2), Ly-6G (PerCP-Cy™5.5; clone 1A8), CD11b (PE-Cy7; clone M1/70)from Biolegend; Siglec-F (PE; clone E50-2440), EpCAM/CD326 (APC; clone 9C4) and SLAMF-6 (PE, clone 13G3) from BD Pharmingen. Monomers for I-A^b^ *Mtb* ESAT6_4-17_, and I-A^b^ Ag85B_280-294_ were kindly provided by the NIH Tetramer Core Facility (https://tetramer.yerkes.emory.edu/). Tetramers were made in house by incubating the monomers with streptavidin-APC from Thermofisher. Cells were fixed and permeabilized using eBioscience Intracellular Fixation and Permeabilization kit (Invitrogen, 88-8824-00). Cell suspensions were then stained with IFN-γ (PerCP/Cy5.5; clone XMG1.2), TNF-α (PE; clone MP6-XT22) from Biolegend IL-2 (BV605; clone JES6-5H4), and TCF-1 (BV421; clone S33-966) from BD Horizon. Cell subsets were defined as follows: Neutrophils (Neut) (Ly-6^Hi^CD11b^Hi^), dendritic cells (DC) (Ly-6G^-/+^MerTK^-^Siglec-F^-^CD11c^+^I-A/I-E^+^), interstitial macrophages (IM) (Ly-6G^/+^MerTK^+^CD64^+^SIglec-F^-^), alveolar macrophages (AM) (Ly-6G^-/+^MerTK^+^CD64^+^SIglec-F^+^), Effector memory T (T_EM_) cells (CD44^+^CD62L^-^CD69^-^CD103^-^), central memory T (T_CM_) cells (CD44^+^CD62L^+^), resident memory T (T_RM_) cells (CD44^+^CD62L^-^CD69^+^CD103^+^), CD69^+^CD103^-^cells (CD44^+^CD62L^-^CD69^+^CD103^-^), stem cell-like memory T cells (TCRβ^+^KLRG1^-^TCF-1^+^)^70^. Samples were acquired on a BD LSRFortessa X-20 cytometer using FACSDiva software (BD Biosciences). The data were analysed by FlowJo software, versions 9 and 10 (Treestar). Absolute cell numbers were calculated using counting beads.

### ELISA

Blood was collected into Z-gel tubes (Sarstedt) and serum was prepared by centrifugation of clotted blood at 8000 rpm for 8 minutes and stored at −80°C until use. ELISA was used to analyse the specific IgG and IgA levels on sera and BALF samples, respectively, for a three epitopes mixture containing Rv3874_76-90_ (CFP10), Rv1886c_280-294_ (Ag85b) and Rv2031c_280-294_ (HspX). Costar 96 well plates (Corning, USA, 3590) were coated with the 3-epitope mixture in a carbonate coating buffer at room temperature for 8 h, then washed with PBST and blocked with 5% skim milk PBST overnight at 4°C. After washing 4 times with PBST, plates were incubated with the diluted samples in 0.5% skim milk PBST buffer for 2 h in a humidified chamber and 5% CO_2_ incubator at 37°C. Plates were washed again and incubated with either 1:3000 diluted biotin goat anti-mouse IgG (in 0.5% skim milk PBST; Sigma SAB4600004-250µl) or 1:1000 diluted biotin goat anti-mouse IgA (BD, 556978), and then incubated for 2 hours at 37°C. For detection, plates were washed and incubated with 1:1000 diluted HRP-conjugated streptavidin (BD, 554066) in 0.5% skim milk PBST buffer for 1 h at 37°C. Washed plates were incubated with TMB (BD, 555214) in the dark at room temperature for 20 minutes. The reaction was stopped by adding 0.18 M H_2_SO_4_, and then the absorbance was measured at λ 450 nm by POLARstar Omega plate reader (BMG Labtech).

### Serum cytokine and chemokine analysis

Frozen serum samples underwent thawing and preparation in accordance with the specifications outlined in the Bio-Plex Pro Mouse Cytokine 23-Plex assay (BioRad). Subsequently, analysis was performed utilizing a MagPix instrument (Luminex). The data analysis and visualization of cytokine levels were in heatmap format (log_2_ concentration).

### Histology

Lung lobes were fixed in 10% formalin, embedded in paraffin, sectioned (4 μm), and stained with H&E. Images were acquired using an Aperio CS2 scanner and analyzed with Aperio ImageScope and ImageJ. Lung damage was quantified as the percentage of tissue area showing dense cellular infiltration.

### Statistics

Data are presented as mean ± SEM. Statistical analyses were performed using GraphPad Prism 8 or 9. Tests used are indicated in figure legends. One-way ANOVA with Tukey’s multiple-comparison test was applied to log-transformed data where appropriate. P < 0.05 was considered statistically significant.

## Results

### LNP mucosal delivery and expression of GFP-mRNA

Effective delivery of mRNA to lung target cells via mucosal administration requires overcoming multiple physiological barriers, including the mucus layer, mucociliary clearance, and uptake by phagocytic cells. To address these challenges, “stealth” lipid nanoparticles (sLNPs) were used for intratracheal (IT) delivery of mRNA. sLNPs are engineered to evade immune recognition and prolong circulation time, largely through incorporation of polyethylene glycol (PEG) lipids. These PEG lipids consist of a hydrophobic lipid anchor attached to a hydrophilic PEG chain that extends into the surrounding aqueous environment, forming a hydrated surface layer around the nanoparticle. This coating helps prevent aggregation and nonspecific protein interactions, thereby improving colloidal stability. Accordingly, sLNPs are serum-stable, protect their nucleic acid cargo from nuclease degradation, and have shown efficient pulmonary delivery of siRNA following intravenous administration^29–31^. In this study, sLNPs composed of DOTAP, DOPE, cholesterol, and C16 PEG-ceramide were prepared in an aqueous sucrose solution. To assess their ability to deliver intact mRNA and support protein expression after mucosal administration, commercially available eGFP mRNA was formulated into sLNPs and administered via the IT route (Fig. 1A). GFP expression in lung cells was evaluated by flow cytometry at 24 h and 48 h post-administration. An increased number of GFP⁺ cells was observed at 48 h, predominantly among dendritic cells (DC) and neutrophils, with a trend toward higher expression in interstitial macrophages (IM) (Fig. 1B, 1C). GFP fluorescence was also assessed in EpCAM⁺ epithelial cells and alveolar macrophages (AM); however, intrinsic autofluorescence in these populations within the GFP channel limited discrimination of true signal from background relative to control groups (Fig. 1D). Collectively, these findings demonstrate that sLNPs can successfully deliver mRNA to lung cells and support protein expression following mucosal administration.

**Figure 1.**
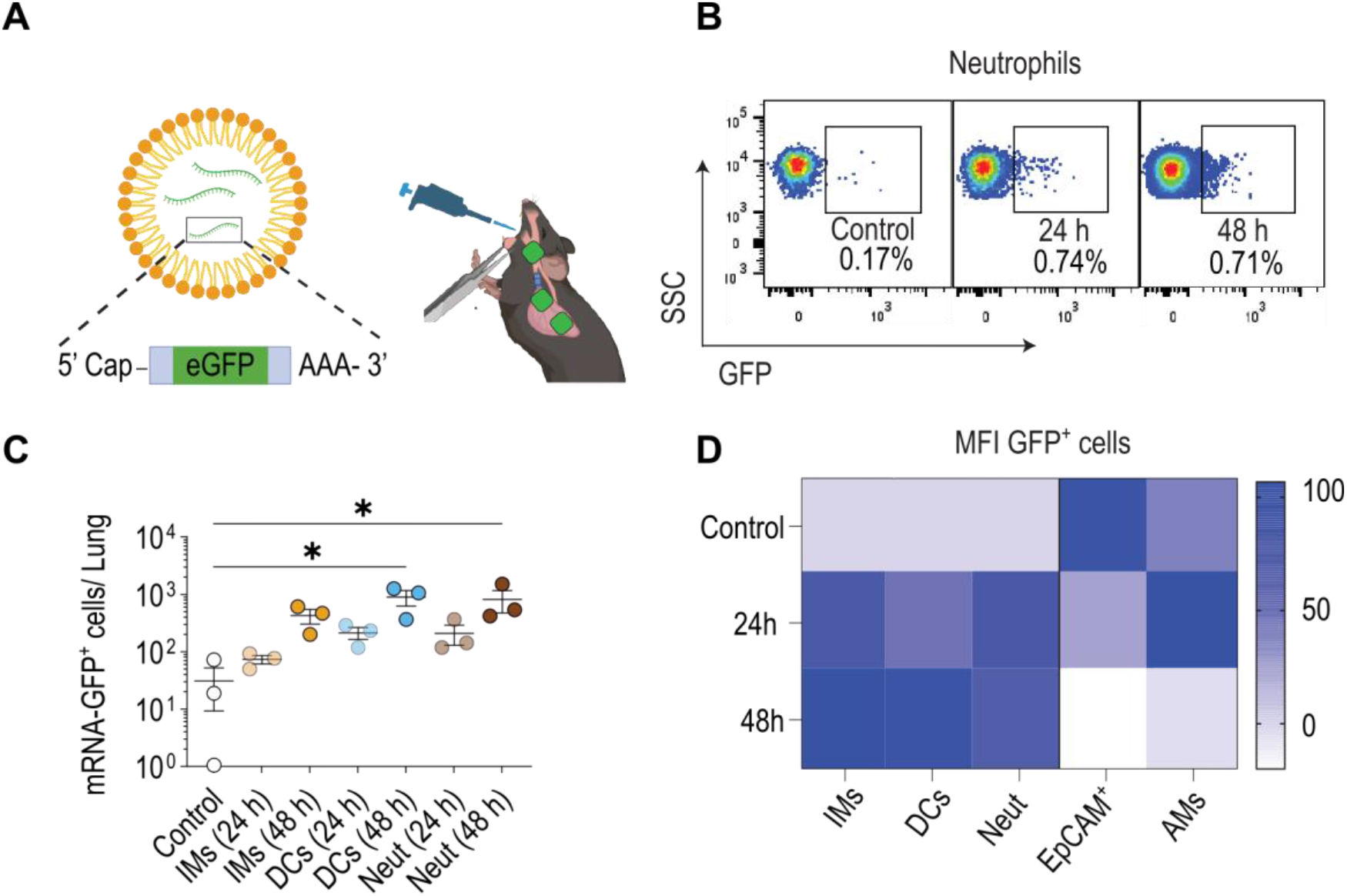
GFP expression by lung cells after intratracheal administration of GFP-mRNA formulated in sLNPs. **A.** Scheme depicting GFP-mRNA formulation in LNPs and IT administration. Lungs were harvest 24 h and 48 h post IT administration with 10 μg of GFP-mRNA-LNPs. Individual cell populations were analysed by flow cytometry for GFP fluorescence. **B.** Concatenated dot plot diagram of GFP^+^ neutrophils. **C.** Number of GFP^+^ cells among IMs, DCs and neutrophils. **D.** Heat map of the mean fluorescence intensity of GFP^+^ cells including EpCAM^+^ cells and AMs. Data was compared by Kruskal-Wallis with Dunn’s multiple comparison post-test. All data show means ± SEM for individual mice. \**p*< 0.05.

### Mucosal delivery of ESAT-6-mRNA enhances antigen-specific T cell responses in BCG-primed mice

We next performed proof-of-concept studies using the immunodominant *Mtb*-derived Early Secreted Antigenic Target 6 (ESAT-6) as a model antigen to characterize antigen-specific immune responses induced by the prime-and-pull vaccination strategy. Customized ESAT-6 mRNA was formulated in sLNPs using the same conditions as for GFP experiments. Mice were primed with recombinant BCG expressing ESAT-6 (BCG::RD1), administered either SC or intratracheally (IT) (Fig. 2A). Prime-and-pull groups received IT ESAT-6–mRNA–sLNPs according to four dosing regimens following SC BCG::RD1 priming: 1) 5 μg once at day 14; 2) 5 μg twice at days 7 and 14; 3) 5 μg three times at days 7, 14, and 21; or 4) a single 10 μg dose at day 14 (Fig. 2B). Antigen-specific T cell responses were analysed 28 days post-vaccination, corresponding to the peak of lung T cell accumulation following mucosal BCG immunization^32^

**Figure 2.**
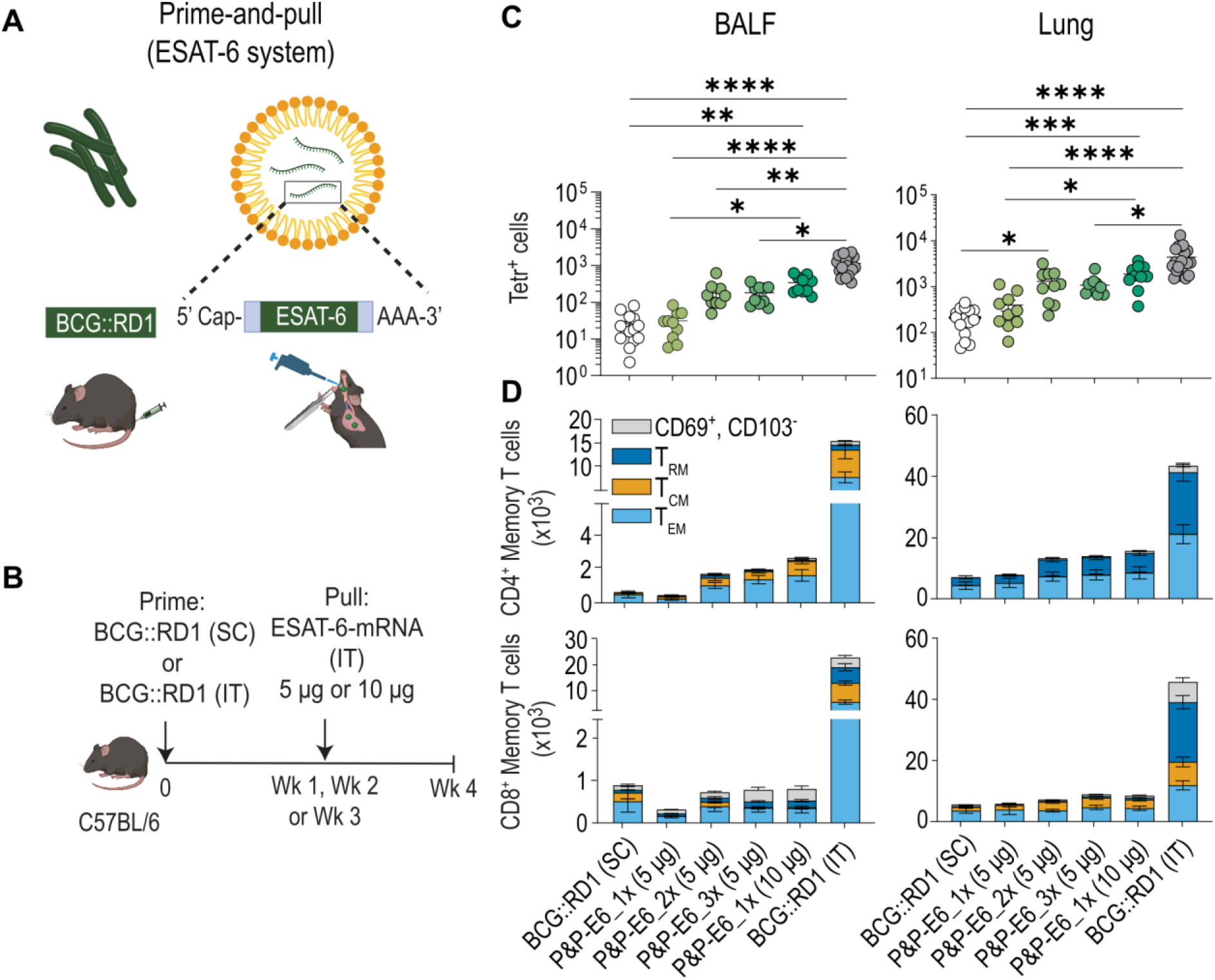
Prime-and-pull vaccination using ESAT-6 as model antigen induces the generation of mucosal specific memory T cells. **A.** Scheme of the prime-and-pull vaccination strategy using BCG::RD1 for priming and ESAT-6-mRNA-sLNPs for the mucosal pulling of immune cells. **B.** C57BL/6 mice were vaccinated with BCG::RD1 either subcutaneously (SC; 1 × 10⁶ CFU) or intratracheally (IT; 1 × 10⁵ CFU). Prime-and-pull groups received IT ESAT-6–mRNA–sLNPs according to four dosing regimens: 1) 5 μg once at day 14; 2) 5 μg twice at days 7 and 14; 3) 5 μg three times at days 7, 14, and 21; or 4) a single 10 μg dose at day 14. Antigen-specific and memory T cell responses were analysed 28 days post-BCG priming. **C.** Number of ESAT-6 tetramer^+^ cells in BALF and lungs. **D.** Number of CD4^+^ and CD8^+^ memory T cell populations (CD69^+^CD103^-^, T_RM_, T_CM_ and T_EM_ cells) in BALF and lungs. Data was compared by Kruskal-Wallis with Dunn’s multiple comparison post-test. NE = 2-3, n = 4 – 5. All data show means ± SEM for individual mice. \**p*< 0.05, \*\**p*< 0.01, \*\*\**p*< 0.001, **** *p*< 0.0001.

A single high-dose administration of ESAT-6 mRNA (10 μg), rather than multiple lower doses, induced significantly higher numbers of ESAT-6 tetramer⁺ T cells in both broncho-alveolar lavage fluid (BALF) and lung tissue compared with SC BCG::RD1 alone (Fig. 2C). As expected, IT BCG::RD1 induced the highest overall numbers of antigen-specific T cells; however, these levels were not significantly greater than those achieved by the single high-dose prime-and-pull regimen. Analysis of CD4⁺ and CD8⁺ memory T cell subsets, including T_EM_, T_CM_, T_RM_, and CD69⁺CD103⁻ populations, revealed a pattern consistent with tetramer⁺ cell frequencies. Notably, a single IT administration of 10 μg ESAT-6 mRNA following SC BCG::RD1 priming significantly increased CD4⁺ memory T cell numbers in the BALF and lungs compared with SC BCG::RD1 alone (Fig. 2D). The number of antigen-specific T and memory T cells increased gradually with every low-dose of ESAT-6 mRNA (i.e 5 μg of the construct). However, a single dose with 5 μg ESAT-6 mRNA induced virtually the same number of cells as SC BCG::RD1. Mucosal vaccination with BCG::RD1 induced the highest accumulation of lung memory T cells, particularly T_EM_ and T_RM_ subsets.

Consistent with systemic priming by SC BCG::RD1, no major differences in antigen-specific or memory T cell numbers were observed in the spleen across vaccination groups (Supp. Fig. 1A, 1C, 1D). Interestingly, IT BCG::RD1 also induced elevated antigen-specific T cell numbers in the spleen, suggesting broader antigen dissemination following mucosal BCG::RD1 administration. In the mediastinal lymph nodes (medLN), only IT BCG::RD1 induced a significant increase in antigen-specific T cells, while the single high-dose prime-and-pull regimen showed a trend toward increased numbers compared with SC BCG::RD1 (Supp. Fig. 1B).

Given the transient nature of lung T cell accumulation, we next assessed responses at day 56 post-vaccination, focusing on SC BCG::RD1, single-dose prime-and-pull, and IT BCG::RD1 groups. Both prime-and-pull vaccination and IT BCG::RD1 induced transient increases in antigen-specific and memory T cells in BALF, lung, medLN, and spleen (Supp. Fig. 2). The most pronounced decline between days 28 and 56 was observed in lungs of IT BCG::RD1-vaccinated mice (Supp. Fig. 2B). Together, these data demonstrate that the prime-and-pull strategy induces robust local memory T cell responses, with a single high-dose mRNA boost achieving responses comparable to IT BCG::RD1 vaccination.

### Immunogenicity of prime-and-pull vaccination targeting multiple candidate antigens

Although BCG::RD1 has demonstrated enhanced protection in murine models of TB compared to conventional BCG, safety concerns in immunocompromised hosts have limited its suitability for human vaccination^33, 34^. We therefore evaluated a prime-and-pull strategy using conventional BCG priming combined with mucosal delivery of mRNA encoding eight candidate proteins, most of which are shared between BCG and *Mtb*. Proteins were selected based on immune reactivity in humans and mice, inclusion of T and B cell epitopes, expression by *Mtb*, and prior evaluation in vaccine candidates (Table 1).

**Table 1.** Selection of candidate proteins and their immunological characteristics.

| Antigen | Function and location | Vaccine candidate | Expression |
| --- | --- | --- | --- |
| **HBHA**<br>(Heparin-binding hemagglutinin adhesin) | Exposed-surface protein involved in adherence to epithelial cells.<sup>71</sup> | Pre-clinical data available.<sup>72</sup> | *Mtb* / BCG |
| **HRP1**<br>(Hypoxic response protein 1, Rv2626c) | Protein present in the cytoplasm, membrane and cell-wall with linked to latency phase of the disease.<sup>73</sup> | MVATG18598 (pre-clinical).<sup>74, 75</sup> | *Mtb* / BCG |
| **HspX** (Rv2031c) | Chaperone protein associated with latency and present in the cytoplasm, membrane and cell wall.<sup>76</sup> | TB/FLU-05E (phase II clinical trials). | *Mtb* / BCG |
| **CFP10** (Rv3874, esxB) | Secreted protein involved in bacterial escape and immune modulation.<sup>76</sup> | AEC/BC02 (phase II), MVATG18598 (pre-clinical), GamTBvac (phase III). | *Mtb* |
| **Ag85b** (Rv1886c) | Cell wall biogenesis and immune response. Secreted protein.<sup>76</sup> | AEC/BC02 (phase II), H56:IC31 (phase IIb), MVATG18598 (pre-clinical), ID91 repRNA (pre-clinical). | *Mtb* / BCG |
| **VapB47** (Rv3407) | Antitoxin for VapC47 toxin. Binds to regulatory DNA sequences.<sup>77</sup> | MVATG18598 (pre-clinical), BCGΔureC::hly (VPM1002) (phase III) BCG AERAS-407 (pre-clinical) | *Mtb* / BCG |
| **PPE18** (Rv1196) | Secreted protein involved in virulence and immune modulation.<sup>76</sup> | M72/AS01E (phase III), MTB72F-AS2A (phase I) | *Mtb* |
| **PE13** (Rv1195) | Secreted protein with potential involvement on bacterial survival within macrophages.<sup>76</sup> | Pre-clinical data available, part of the TCR Informed TB Antigen (TITAN) mRNA vaccine candidate.<sup>78</sup> | *Mtb* / BCG |

Four customized multi-antigen mRNA constructs were generated, each encoding two proteins separated by a glycine-rich linker and preceded by a human IgE signal peptide. All constructs included standard mRNA modifications and were formulated at equal weight ratios into sLNPs (Fig. 3A).

**Figure 3.**
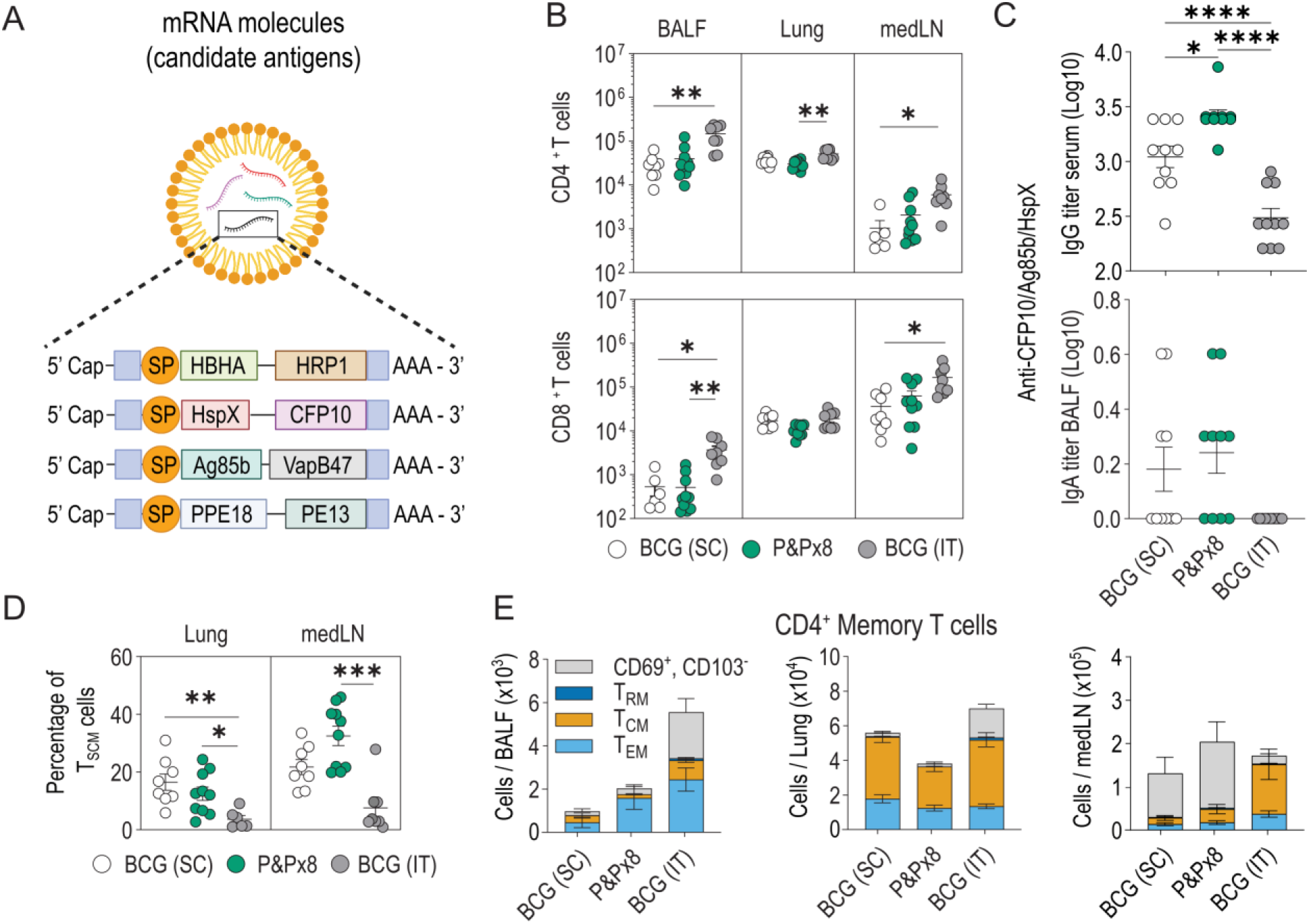
Prime-and-pull targeting candidate antigens induces the generation of respiratory T cells with a stem-like profile. **A.** Scheme of the four mRNA constructs containing 8 different antigens and the formulation in the sLNPs. Hybrid F1 C57BL/6 × BALB/c mice were vaccinated with BCG either SC or IT as previously described. Prime-and-pull groups received IT a single 10 μg dose of mRNA-sLNPs mix-constructs at day 14. T cell populations were analysed 28 days post-BCG priming. **B.** Number of CD4^+^ and CD8^+^ cells in BALF, lungs and medLN. **C.** Titres of CDF-10/Ag85b/HspX epitopes-specific immunoglobulin G (IgG) and IgA in sera and BALF, respectively. **D.** Percentage of T_SCM_ cells in lung and medLN. **D.** Number of CD4^+^ memory T cell populations (CD69^+^CD103^-^, T_RM_, T_CM_ and T_EM_ cells) in BALF, lung and medLN. Data was compared by Kruskal-Wallis with Dunn’s multiple comparison post-test. NE=2, n=4–5. All data show means ± SEM for individual mice. \*\**p*< 0.05, \*\**p*< 0.01, \*\*\**p*< 0.001, **** *p*< 0.0001.

To broaden the major histocompatibility complex (MHC) profile, hybrid F1 C57BL/6 × BALB/c mice were used. Flow cytometric analysis showed that prime-and-pull vaccination targeting the candidate antigens induced comparable numbers of total CD4⁺, CD8⁺, and CD4⁺ memory T cells in BALF, lung, and medLN relative to SC BCG vaccination (Fig. 3B, 3E). As expected, IT BCG induced higher local T cell numbers than systemic vaccination. Further analysis using MHC-II Ag85b tetramers revealed similar numbers of Ag85b-specific T cells across groups, with IT BCG inducing the highest numbers (Supp. Fig. 3A). No significant differences were observed in splenic T cell populations (Supp. Fig. 3B, 3C). However, because tetramer reagents were available only for Ag85b, the antigenic specificity of the broader cellular response could not be resolved across all encoded proteins.

Serum antibody analysis showed that prime-and-pull vaccination significantly increased IgG titers against CFP-10, Ag85b, and HspX compared with both SC and IT BCG groups, indicating enhanced humoral immunity. It is important to note that CFP-10 is absent from BCG, and therefore comparisons of CFP-10-specific responses between BCG and CFP-10-containing vaccine candidates should be interpreted with caution.

Stem cell-like memory T (T_SCM_) cells are key drivers of durable T cell immunity and have been implicated in the efficacy of highly effective vaccines. T_SCM_ are characterised by long-term persistence, strong proliferative capacity, and the ability to generate effector T cells upon antigen re-exposure^35–40^. Given the short-lived nature of lung memory T cells induced by both prime-and-pull vaccination and IT BCG, we examined T_SCM_ cell frequencies after immunisation. Prime-and-pull vaccination was associated with higher T_SCM_ cell frequencies in the lung and medLN compared with IT BCG (Fig. 3D).

Collectively, these data indicate that prime-and-pull vaccination induced cellular responses comparable to SC BCG in BALF, lung, and medLN, while also enhancing antibody responses and showing higher T_SCM_ frequencies than IT BCG.

### Prime-and-pull vaccination confers protection comparable to mucosal BCG

To evaluate protective efficacy, mice vaccinated with SC BCG, prime-and-pull, or IT BCG were challenged with aerosolized *Mtb* 60 days post-immunization. Two independent studies were conducted using mRNA-sLNPs targeting either ESAT-6 or the eight candidate antigens. Mirroring the intermediate numbers of T cells (Fig. 2C, D), in the ESAT-6 model, prime-and-pull vaccination also conferred an intermediate level of protection, with reduced lung bacterial burden and pathology compared with SC BCG::RD1 but lower protection than IT BCG::RD1 (Fig. 4A–C).

**Figure 4.**
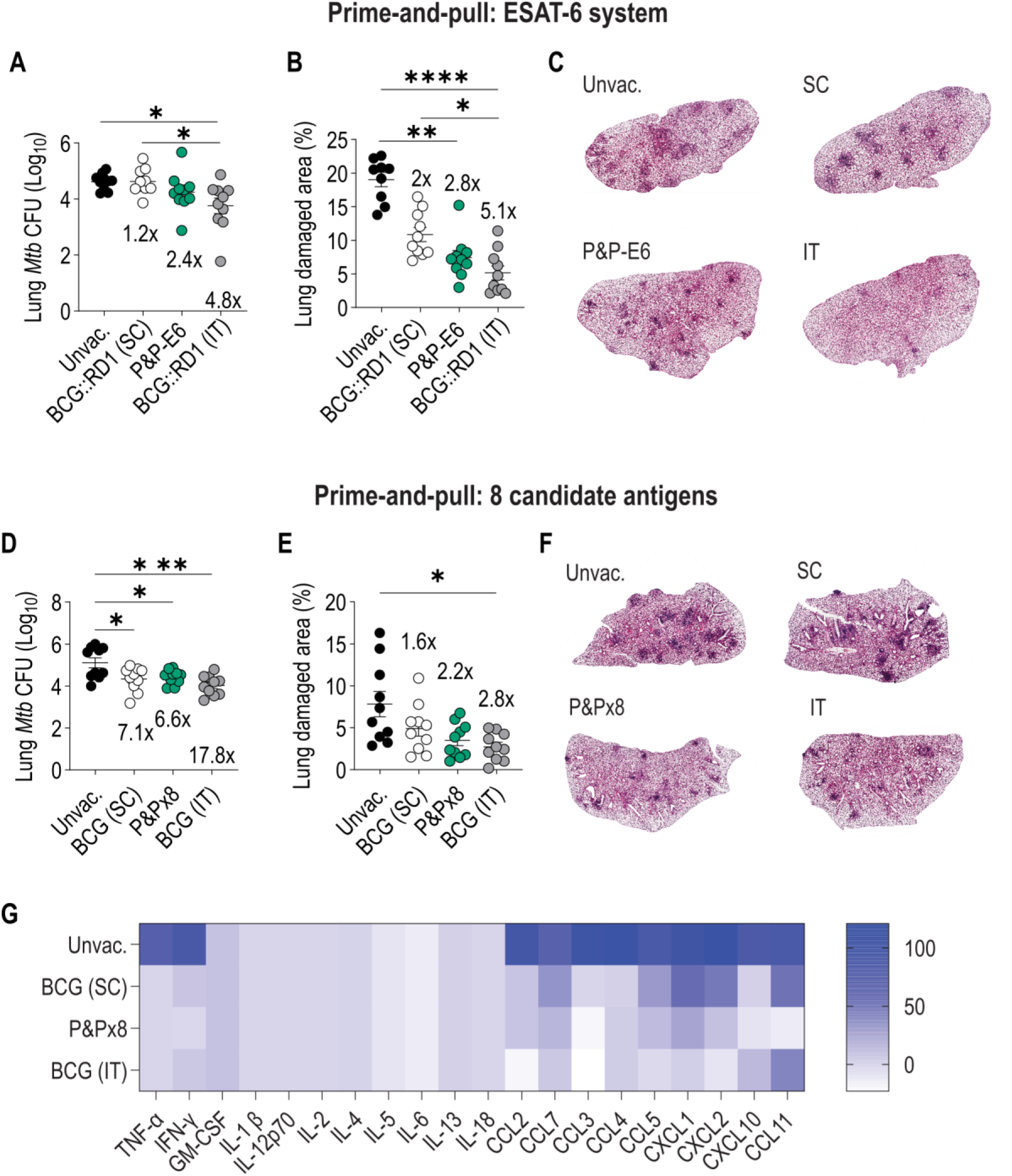
Prime-and-pull vaccination induces similar protection levels against *Mtb* challenge compared to mucosal BCG vaccination. In A. –. **C.** C57BL/6 mice were vaccinated with BCG::RD1 either SC or IT; prime-and-pull groups received IT a single 10 μg dose of at day 14. In **D. – G.** Hybrid F1 C57BL/6 × BALB/c mice were vaccinated with BCG either SC or IT, and prime-and-pull groups received IT a single 10 μg dose of mRNA-sLNPs mix-constructs at day 14. Sixty days later, mice were aerosol infected with very low dose of *Mtb* H37Rv. *Mtb* bacterial burden was determined 45 days post infection in lungs and spleen. Lung damage was determined as the percentage of the area of dense cell infiltration in H&E-stained lung sections. ESAT-6 system: **A.** Bacterial burden in lung. **B.** Percentrage of lung damaged area. **C.** Representative images of lung infiltration. 8 candidate antigens: **D.** Bacterial burden in lung. **E.** Percentrage of lung damaged area. **F.** Representative images of lung infiltration. **G.** Profile of cytokines and chemokines in serum. For the bacterial burden, data was log-transformed and differences tested using the one-way ANOVA with Tukey’s multiple comparison. NE = 2, n = 5. All data show means ± SEM or individual mice. \**p*< 0.05, \*\**p*< 0.01, \*\*\**p*< 0.001, **** *p*< 0.0001. Fold-change was calculated as the ratio of median of the raw data per each vaccination group over the unvaccinated group.

In studies targeting the eight candidate antigens, hybrid F1 mice exhibited higher lung bacterial burdens overall compared with C57BL/6 mice, but reduced lung pathology following infection (Fig. 4). All vaccination groups significantly reduced lung bacterial loads compared with unvaccinated controls, with comparable reductions observed between SC BCG and prime-and-pull groups (Fig. 4D). IT BCG induced the greatest reduction (∼18-fold). Differences in lung pathology were less pronounced, with significant reductions observed only in IT BCG-vaccinated mice (Fig. 4E, 4F).

Elevated pro-inflammatory mediators in unvaccinated mice indicated an active infection (Fig. 4G). While SC BCG partially reduced inflammatory signatures, both prime-and-pull and IT BCG induced similarly low levels of inflammatory mediators. These findings indicate that prime-and-pull vaccination confers protection comparable to IT BCG while limiting systemic inflammation.

### Prime-and-pull vaccination provides a sweet spot for inflammation, immunogenicity and efficacy

Finally, we directly compared inflammatory, immunogenic, and protective parameters across vaccination strategies. In the ESAT-6 antigen system, IT BCG induced the highest numbers of polyfunctional ESAT-6-specific IFN-γ⁺TNF-α⁺IL-2⁺ CD4⁺ and CD8⁺ T cells in the lung, whereas prime-and-pull vaccination elicited lower, but still appreciable, responses that were comparable to those induced by SC BCG (Fig. 5A). This was accompanied by significantly greater lung pathology at 4 weeks post-vaccination, particularly in the BCG::RD1 group, whereas prime-and-pull and SC BCG were associated with similarly low levels of lung damage (Fig. 5B). In the eight-antigen system, a similar pattern was observed, with prime-and-pull vaccination again achieving an intermediate pro-inflammatory cytokine profile and reduced lung damage after vaccination relative to SC and IT BCG.

**Figure 5.**
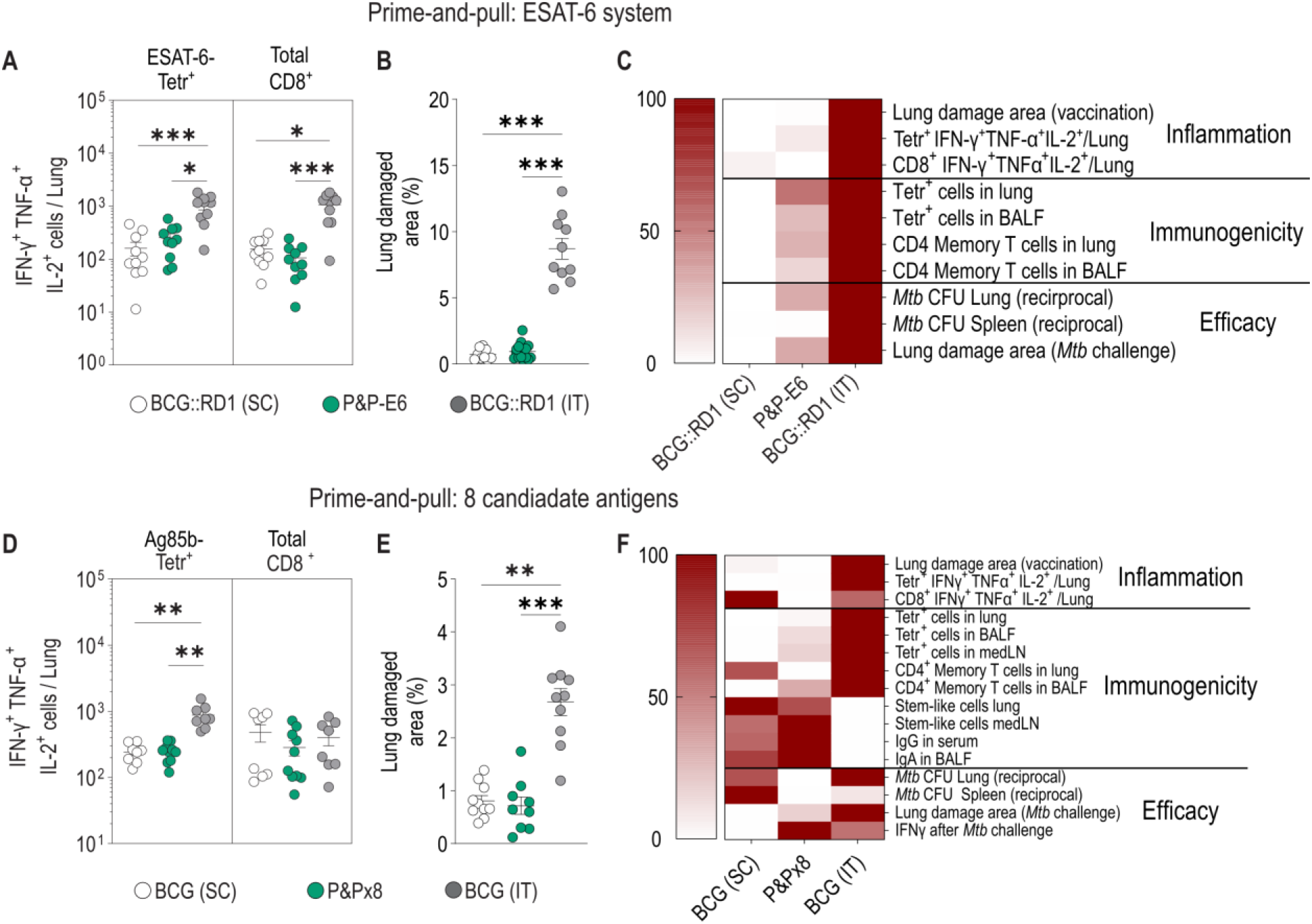
Prime-and-pull vaccination induces similar immunogenicity and protection compared to mucosal BCG vaccination with a reduced pro-inflammatory profile. A. –. **C.** Results from the ESAT-6 system and **D. – F.** Data from the 8 candidate antigens study. Mice were vaccinated as previously described in Fig. 2 and Fig. 3, respectively. Cytokine producing cells and lung cell infiltrations were analysed 28 days post priming. ESAT-6 system: **A.** Number of lung ESAT-6-tetramer^+^ and total CD8^+^ cells producing IFN-γ, TNF-α and IL-2⁺. **B.** Percentrage of lung damaged area at 4 weeks post-vaccination. **C.** Heat map comparing several inflammatory, immunological, and efficacy parameters. For the efficacy data, reciprocal values were used. Data were normalized, setting the smallest value to 0 and the largest to 100 and the heat map was constructed based on the median. 8 candidate antigens: **D.** Number of lung Ag85b-tetramer^+^ and total CD8^+^ cells producing IFN-γ, TNF-α and IL-2⁺. **E.** Percentrage of lung damaged area at 4 weeks-post-vaccination. **F.** Heat map comparing several inflammatory, immunological, and efficacy parameters. Data was compared by Kruskal-Wallis with Dunn’s multiple comparison post-test. NE = 2 - 3, n = 4 – 5. All data show means ± SEM for individual mice. \**p*< 0.05, \*\**p*< 0.01, \*\*\**p*< 0.001.

The integrated heatmap analysis summarized inflammatory, immunological, and protective readouts across the two antigen systems. In both cases, prime-and-pull vaccination clustered as a balanced regimen, with low inflammation, intermediate immunogenicity, and substantial protective efficacy relative to the comparison groups (Fig. 5C, 5F). In the ESAT-6 system, this profile was reflected by reduced inflammatory burden alongside preserved antigen-specific T-cell responses and protection. In the eight-antigen system, a similar pattern was observed, with prime-and-pull maintaining favourable efficacy while avoiding the higher inflammatory profile associated with IT BCG. Collectively, these data support mRNA-based prime-and-pull vaccination as a balanced strategy that limits excessive inflammation while maintaining immunogenicity and protection.

## Discussion

The development of improved vaccines and alternative vaccination strategies remains central to achieving TB reduction, elimination, and potentially eradication in line with WHO goals. In this study, we show that a prime-and-pull vaccination strategy can promote local pulmonary immunity while limiting the inflammatory burden associated with conventional mucosal vaccination. By combining systemic BCG priming with mucosal mRNA delivery, this approach recruited antigen-specific immune responses to the lung and mediastinal lymph node while causing substantially less histopathological damage than intratracheal BCG vaccination. Together, these findings support prime-and-pull vaccination as a proof-of-principle platform for respiratory TB immunization.

A central feature of this strategy is its ability to uncouple immune recruitment from tissue-destructive inflammation. The lung is uniquely vulnerable to immunopathology, and respiratory vaccines must therefore induce sufficient inflammation to drive immune activation without causing excessive tissue damage. Maintaining a level of inflammation that is low enough to preserve tissue integrity yet sufficient to prime protective immunity is critical for effective pulmonary vaccination^41, 42^. Accumulating evidence shows that chronic inflammation can impair vaccine efficacy and contribute to severe pathological outcomes^43–46^. In the airways, dysregulated inflammation can result in alveolar damage, impaired blood oxygenation, respiratory distress, and long-term sequelae such as lung fibrosis^47^. Consistent with this, we previously showed that mucosal vaccination with conventional BCG and BCG::RD1 induces moderate and severe lung inflammation, respectively^48^. This pathology was characterised by haemorrhagic lesions, peribronchiolar and perivascular immune cell infiltrates, and the presence of interstitial giant cells and foamy macrophages^48^. Our data extend these observations by showing that prime-and-pull vaccination achieved local immune recruitment with substantially less lung pathology than intratracheal BCG.

The use of sLNPs for mucosal delivery is a key component of this platform. Most mRNA vaccines, including recent COVID-19 formulations, were optimised for intramuscular administration. For example, while macaques vaccinated with a COVID-19 mRNA vaccine via the intramuscular route developed highly protective immune responses, intranasal boosting with the same vaccine failed to enhance immunogenicity or efficacy^49, 50^. Notably, in that same study, intratracheal boosting with an adenovirus-based vaccine led to near-complete protection against high-dose mucosal SARS-CoV-2 challenge^50^. Similarly, murine studies using clinically approved lipids showed that although intranasal administration of mRNA-LNPs produced high transfection efficiency and protein expression in vivo, immune responses remained limited^51^. These findings underscore the need for improved formulations for effective delivery of mRNA vaccines to the airways. Here, we show that sLNPs not only support delivery and translation of encapsulated cargo but also promote increased numbers of endogenous antigen-specific T and memory cells in the lungs of prime-and-pull-vaccinated mice. This is consistent with previous studies showing that intranasal mRNA-LNP boosters can enhance lung immunity in animals previously primed systemically^51, 52^.

An additional strength of this strategy is the induction of stem cell-like memory T cells in the lung and mediastinal lymph node. The limited persistence of lung T_RM_ cells remains a major challenge for respiratory vaccines aimed at generating durable T cell-mediated immunity in the airway mucosa. T_SCM_ cells have emerged as an important source of effector memory T-cell replenishment upon antigenic recall^40^. These cells accumulate in lymph nodes and express markers such as TCF-1, which supports self-renewal and long-term survival^38, 40^. Because the medLN drain the respiratory tract, they may serve as a reservoir for this stem-like population over extended periods. In this study, we observed an increased percentage of T_SCM_ cells in the medLN of prime-and-pull-vaccinated mice compared with those receiving mucosal BCG. These findings are consistent with the potential for more durable immune maintenance, although long-term longitudinal studies will be needed to confirm persistence over time.

Our results also showed that the multivalent mRNA approach broadened humoral responses. Serum antibody analysis showed that prime-and-pull vaccination significantly increased IgG titres against CFP-10, Ag85b, and HspX compared with both SC and IT BCG groups, indicating enhanced humoral immunity. More broadly, the multivalent design is consistent with the antigen-selection criteria used here, namely immune reactivity in humans and mice, inclusion of T and B cell epitopes, expression by *Mtb*, and prior evaluation in vaccine candidates. This matters because the antigenic breadth of the construct may improve the likelihood of coverage across diverse human MHC backgrounds. In principle, full-length antigen-encoding mRNA should generate multiple processed peptides that can be presented across a broader range of HLA class I and II molecules than epitope-restricted approaches, thereby increasing the chance of population-level coverage.

The multivalent nature of the vaccine construct should therefore be viewed as a strategy for breadth and feasibility rather than as a final universal formulation. The eight antigens included in this study were selected on the basis of prior immunological evidence and antigenic relevance, but the optimal antigen combination for broad human protection remains unresolved. Further optimisation of antigen composition may still be required to identify the most effective formulation for broad human protection and to surpass the protection levels currently set by mucosal live-attenuated vaccines. In this sense, the present work demonstrates the platform potential of prime-and-pull vaccination rather than claiming that the exact construct tested here is ready for direct clinical deployment.

The translational path for a prime-and-pull strategy will also require careful consideration of delivery. Here, intratracheal administration was used to ensure controlled dosing and reproducibility in mice. For human application, however, aerosolized or intranasal delivery would be more practical than direct intratracheal administration. Mucosal vaccination is already an active area of clinical development, and the broader feasibility of respiratory immunization is supported by ongoing intranasal vaccine studies^53, 54^. Nonetheless, successful translation will depend on optimising formulation stability, lung deposition, mucociliary clearance, and tolerability in humans. Thus, the most realistic near-term goal is iterative refinement of a mucosal platform that can be adapted for human use.

The prime-and-pull strategy also builds on the established benefits of BCG vaccination. It is well documented that BCG confers beneficial non-specific effects and reduces mortality from other infections. Meta-analyses from three randomized trials in Guinea-Bissau, including over 6,500 newborns, showed that early BCG vaccination reduced infant (≤12 months) and neonatal (≤28 days) mortality by 16% and 38%, respectively^55–57^. Another randomized trial in India found that early vaccination with BCG and oral polio vaccine reduced all-cause neonatal mortality by 17% and infection-related mortality by 47%^58^. These effects likely reflect BCG’s capacity to induce trained immunity through epigenetic reprogramming in monocytes, neutrophils, macrophages, and NK cells^59^. This state is characterised by an increased capacity to secrete pro-inflammatory cytokines, leading to non-specific protection against malaria, sepsis, and respiratory infections^60–62^. Consequently, maintaining systemic BCG regimens in countries with high infectious disease burdens remains essential for preventing childhood mortality. Our prime-and-pull candidate builds on this established regimen by preserving the benefits of systemic BCG while adding a targeted respiratory immune response aimed specifically at pulmonary TB, the most prevalent and transmissible form of the disease.

Recent investigations have shown that selective organ targeting of therapeutic cargo is possible by altering the internal or external charge of LNPs^63^. Tuning the proportion of quaternary ammonium lipids, such as DOTAP, can endow LNPs with a distinctive protein corona that promotes lung-specific mRNA delivery after intravenous administration^63–65^. The surface of LNPs can also be modified with antibodies, peptides, or sugars for targeted delivery. For example, systemic administration of mRNA-LNPs conjugated with antibodies specific to vascular cell adhesion molecule (PECAM-1) resulted in markedly increased protein production in the lung^66^. Such advances in targeted mRNA delivery are highly relevant to vaccine design because they may reduce dose requirements and minimize off-target effects. In our system, the use of sLNPs likely contributed to the favourable balance between local pulmonary delivery and reduced inflammatory toxicity, supporting the broader feasibility of rationally engineered lung-targeted nucleic acid vaccines. Our findings further suggest that antigen-specific T-cell responses may be dose-dependent and subject to a threshold effect, as a single 10 μg dose outperformed multiple smaller doses of mRNA-sLNPs. This is consistent with evidence that innate sensing and type I IFN responses induced by LNPs can blunt responses to repeated dosing.^67, 68^. In the multivalent prime-and-pull setting, the mRNA dose per antigen was substantially lower than in the single-antigen ESAT-6 formulation, which likely contributed to the more modest differences observed relative to SC BCG. Future optimisation may therefore benefit from dose-sparing strategies, such as self-amplifying RNA, to sustain antigen expression at lower input amounts^64, 66, 67^.

While these findings support prime-and-pull as a feasible vaccination strategy, several important limitations should be considered. First, although we observed significant induction of T_SCM_ cells in the lungs and medLN, protective efficacy was evaluated primarily at 60 days post-vaccination, and longer longitudinal studies will be needed to determine whether these stem-like populations sustain durable protection over extended periods. This is particularly relevant given the relatively transient nature of lung T-cell responses. Second, intratracheal administration was used here to ensure precise and reproducible dosing in mice; however, clinical translation will likely require aerosolized or intranasal delivery^23, 28, 54^. These alternative mucosal routes introduce additional physiological variables, such as deposition patterns and mucociliary clearance, which may influence sLNP immunogenicity. Third, although the hybrid F1 model enabled broader MHC profiling, murine systems cannot fully recapitulate the complexity of human tuberculosis, including granulomatous pathology, latency, and reactivation^69^. Finally, while the multivalent mRNA constructs targeted eight immunodominant proteins, further refinement of antigen composition may still be required to identify the most effective formulation for broad human protection.

In conclusion, our study shows that the prime-and-pull strategy can direct local pulmonary immunity while limiting the inflammatory burden associated with live mucosal vaccination. By combining systemic BCG priming with mucosal mRNA delivery, this platform enabled antigen-specific immune responses in the lung and supported protective efficacy with reduced tissue pathology. Together, these findings establish prime-and-pull vaccination as a credible and adaptable framework for further optimisation in respiratory TB vaccine development.

## Data availability statement

All data are available in the main text or the supplementary materials. Requests for further information and resources should be directed to and will be fulfilled by the lead contact.

## Acknowledgements

We acknowledge the NanoMedicine Biofoundry (Institute for Biomedicine and Glycomics, Griffith University) for the design, formulation, and supply of the LNP materials used. We also acknowledge the NIH Tetramer Core Facility for kindly providing monomers and instructions for tetramerization. The laboratory of Andreas Kupz is supported by an NHMRC Investigator Grant (APP2008715). The funder played no role in study design, data collection, analysis and interpretation of data, or the writing of this manuscript.

## Author contribution

Conceptualization: AK, AMVH. Methodology: AK, AMVH. Investigation: AMVH, GZ, JS, HDS, SMH, MP. Manuscript writing: AMVH, AK. Funding acquisition: AK, AMVH.

## Supplementary Information

**Supplementary Figure 1.**
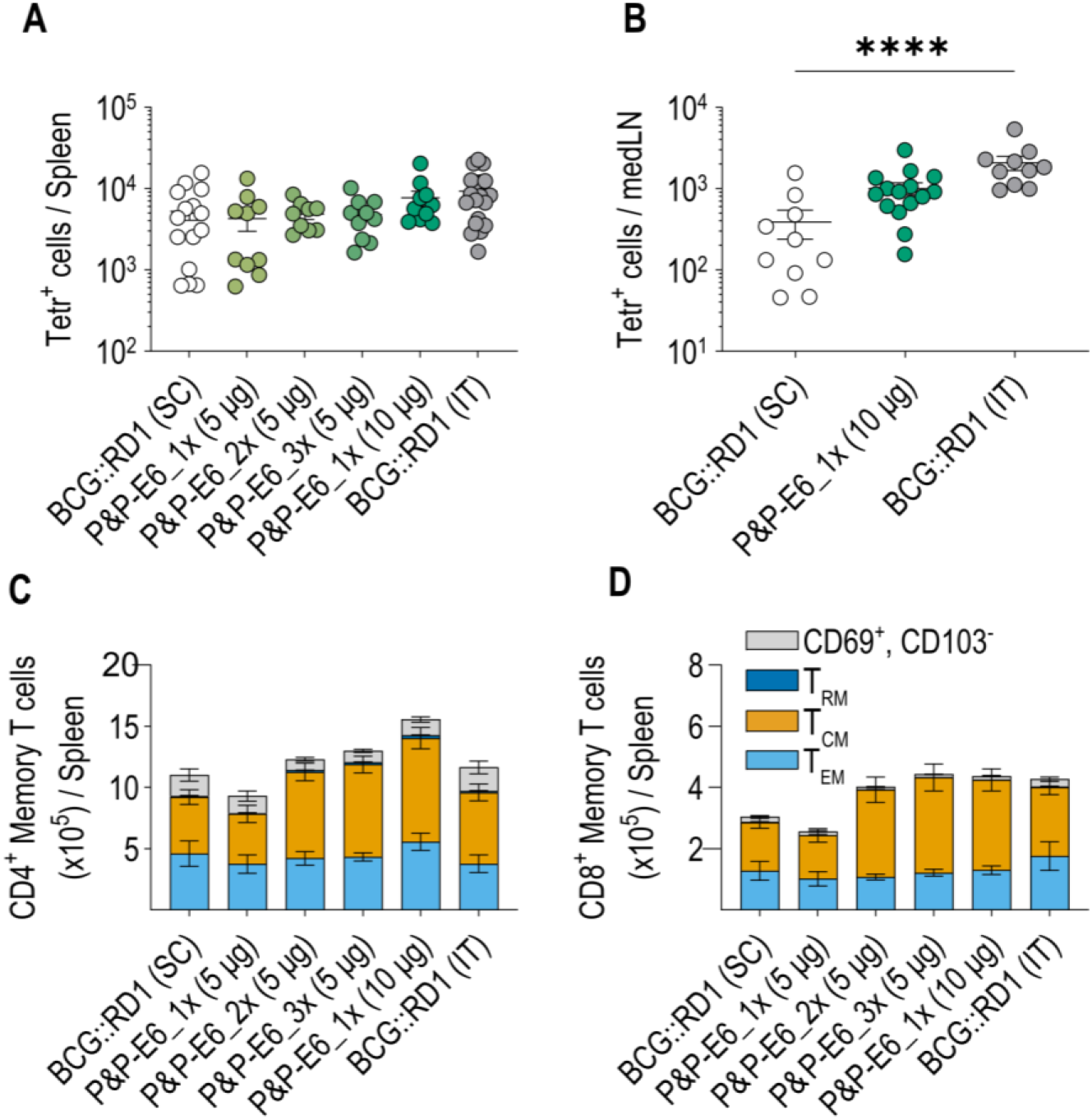
Prime-and-pull vaccination targeting ESAT-6 as model antigen induces systemic T cells responses. C57BL/6 mice were vaccinated as described in Fig. 2. Antigen-specific and memory T cell responses were analysed 28 days post-BCG priming. **A.** Number of ESAT-6 tetramer^+^ cells in spleen. **B.** Number of ESAT-6 tetramer^+^ cells in medLN. **C.** and **D.** Number of CD4^+^ and CD8^+^ memory T cell populations (CD69^+^CD103*, **T_RM_,** T_CM_ and T_EM_ cells) in spleen. Data was compared by Kruskal-Wallis with Dunn’s multiple comparison post-test. NE = 2-3, n = 4 - 5. All data show means ± SEM for individual mice. \*\*\*\**p< 0.0001*.

**Supplementary Figure 2.**
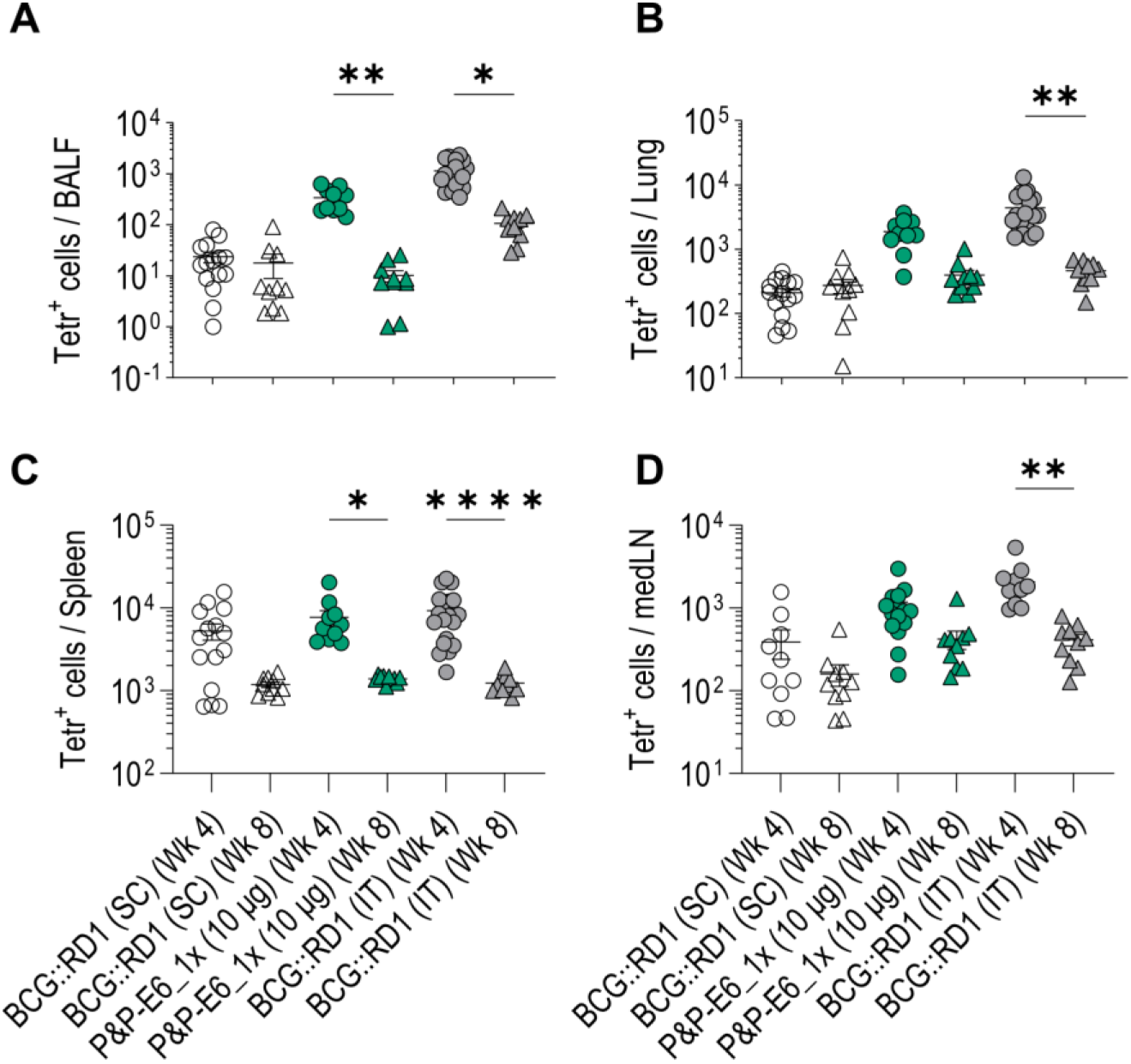
Mucosal vaccination with BCG::RD1 or prime-and-pull induces a transient respiratory T cell response. C57BL/6 mice were vaccinated as described in Fig. 2. Antigen-specific and memory T cell responses were analysed at 28- and 56-days post-BCG priming. **A., B., C.,** and **D.** Number of ESAT-6 tetramer* cells in BALF, lung, spleen and medLN, respectively. Data was compared by Kruskal-Wallis with Dunn’s multiple comparison post-test. NE = 2-3, n = 4-5. All data show means ± SEM for individual mice. *\*p< 0.05, **p< 0.01, ***p< 0.001, ****p< 0.001,*.

**Supplementary Figure 3.**
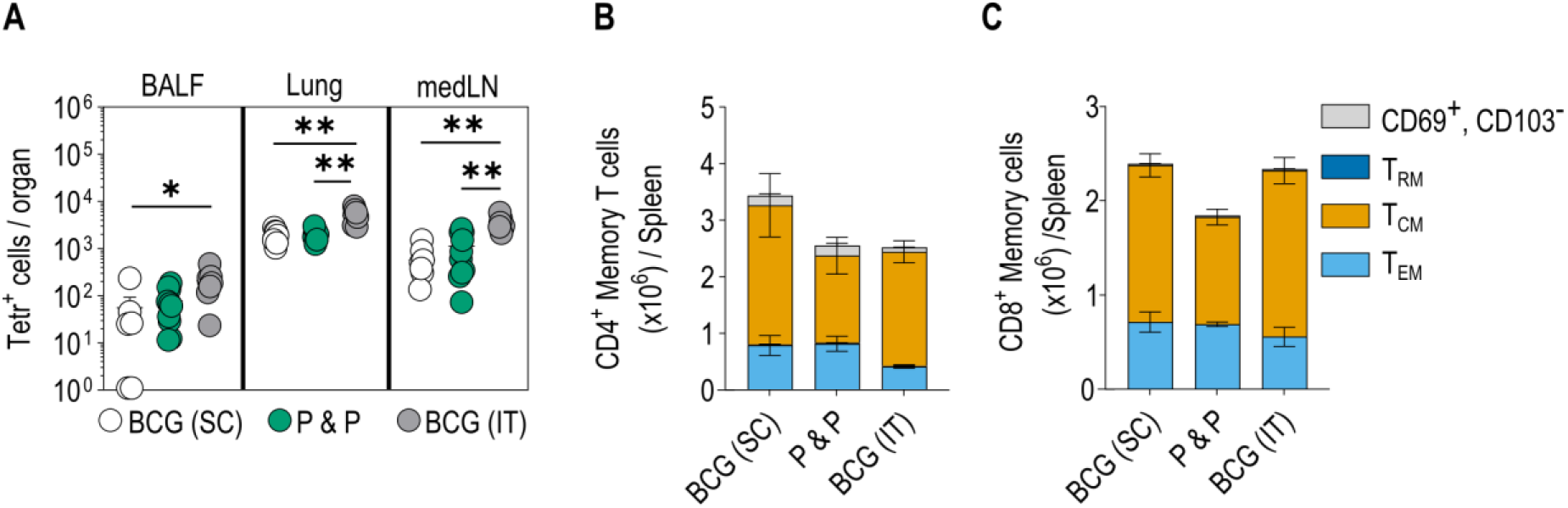
Antigen specific T cell responses to prime-and-pull vaccination targeting 8 candidate antigens. Hybrid F1 C57BL/6 x BALB/c mice were vaccinated as described in Fig. 3. Antigen-specific and memory **T** cell responses were analysed 28-days post-BCG priming. **A.** Number of Ag85b tetramer^+^ cells in BALF, lung, and medLN. B. and **C.** Number of CD4^+^ and CD8^+^ memory **T** cells in spleen, respectively. Data was compared by Kruskal-Wallis with Dunn’s multiple comparison post-test. NE = 2-3, **n** = 4 - 5. All data show means ± SEM for individual mice. *\*p< 0.05, **p< 0.01, ***p< 0.001, **** p<0.0001*.

